# IgG Propels Atherosclerosis by Noncanonically Activating Macrophages

**DOI:** 10.64898/2026.03.13.711718

**Authors:** Tarik Zahr, Kai Zhang, Shengkang Huang, Chenyi Xue, Lexiang Yu, Zhewei Yu, Bing Li, Dongyang Liang, Qian Wang, Mariya Shadrina, Qianfen Wan, Xueming Li, Fuping You, Amy R. Kontorovich, Muredach P. Reilly, Zhan Hu, Wei Feng, Liheng Wang, Li Qiang

**Author notes:** These authors contributed equally to this work (Tarik Zahr, Kai Zhang, Shengkang Huang).

## Abstract

Despite being a central component of adaptive immunity and a highly abundant serum protein, the contribution of IgG to the milieu of atherosclerosis remains unappreciated. Here, we identify a pro-atherogenic role for IgG as it activates an innate immune cascade, independent of its classical antigen-neutralizing function. Analyses of human coronary artery plaques reveal a positive correlation between IgG and cardiovascular and cerebrovascular disease severity. Integrated single-cell plaque analyses localize IgG, coinciding with its recycling receptor FcRn, to pro-inflammatory and foamy macrophages. Genetic ablation of FcRn in myeloid cells prevents IgG from accumulating in mouse atherosclerotic lesions, diminishing plaque size and inflammation. Mechanistically, IgG acts as an endogenous ligand for TLR4, triggering NF-κB–NLRP3 inflammasome signaling without requiring its antigen-binding domain. Additionally, IgG accelerates macrophage foam cell formation through upregulation of downstream effector LCN2. Our work uncovers a role for previously overlooked adaptive immune molecules in the pathogenesis of atherosclerosis through a noncanonical mechanism linked with innate immunity.

## INTRODUCTION

Cardiovascular diseases (CVD) are the leading cause of global mortality, a burden primarily driven by atherosclerosis and its complications^1^. This multifactorial disease is characterized by fatty streak formation along the arterial wall^2^, progressing as elevated circulating low-density lipoproteins (LDL) accumulate in the subendothelial intima^3^. Following oxidative modifications, oxLDL is internalized by macrophages, leading to foam cell formation^4^. This prolonged accumulation develops into a necrotic core, exacerbating plaque instability and rupture risk^5^. Despite effective management of hypercholesterolemia by lipid-lowering drugs, only one-third of CVD risk is controlled^6,7^. This persistent residual risk underscores the urgent need to deepen our understanding of mechanisms of atherogenesis and to identify novel pathogenic factors for improved therapeutic and preventative strategies.

The immune system, comprising of innate and adaptive branches, is essential for host defense and plays a critical role in the pathogenesis of diseases like atherosclerosis ^8,9^. The classical paradigm holds that the innate immune system provides a rapid, non-specific first line of defense, which is subsequently refined by the more specialized adaptive response^10^. In atherosclerosis, a key research challenge is identifying the endogenous damage-associated molecular patterns (DAMPs) that trigger innate immune responses occurring in the arterial wall, as there is little evidence implicating pathogens as the primary inducers^9,11^. These innate responses are not isolated; antigen-presenting cells of innate origin shape subsequent adaptive responses within atherosclerotic lesions ^12,13^, and innate-derived cytokines (e.g., IL-1, TNF-α, and IFN) promote inflammation and activate adaptive immune cells, ensuring a coordinated defense^14^. While substantial evidence supports the classic view that innate immunity directs the adaptive response, this perspective invites a crucial reciprocal question: can adaptive immune components, in turn, regulate innate immunity and influence atherosclerosis progression?

Atherosclerotic plaques are continuously exposed to circulating proteins, including immunoglobulins (Igs). Immunoglobulin G (IgG), the most abundant antibody (∼80% of serum Igs), comprises antigen-binding (Fab) and crystallizable fragment (Fc) regions. Traditionally, IgG is thought to contribute to atherosclerosis via the formation of immune complexes (ICs) with antigens like oxLDL^15^. These ICs engage with Fc gamma receptors (FcγRs) to drive pro-inflammatory responses^16^. IC-activated macrophages exhibit gene expression profiles resembling those in vulnerable carotid plaques^17^, and clinical studies report positive correlations between oxLDL-IC levels and CVD severity. While antigen-specific autoantibodies contribute to atherosclerosis development^26^, they are unlikely to fully account for the strong association between total IgG levels and CVD risk^18–21^. In contrast to the less abundant IgM that has gained more attention in atherosclerotic plaques^25^, the role of IgG per se in atherosclerosis has remained overlooked. Recently, IgG has been implicated in aging, obesity, and metabolic dysfunction^22–24^. Given that aging and obesity are two major atherosclerosis accelerants, we hypothesized that IgG may directly contribute to atherosclerotic disease progression through mechanisms beyond classical antigen recognition.

Moving beyond its traditional role in adaptive immunity, we position IgG as a key component of atherosclerosis, particularly within the plaque microenvironment. Integrating clinical data, genetic approaches and mechanistic studies, we reveal a noncanonical mechanism for IgG to act through the TLR4/NF-κB axis, to stimulate inflammasome signaling and lipocalin–2 (LCN2)-mediated macrophage foam cell formation, independent of its antigen-binding Fab domain. Collectively, these findings uncover a pathogenic role for IgG in atherosclerosis through a mechanism that directly bridges adaptive and innate immunity, pointing to novel avenues for therapeutic intervention.

## RESULTS

### Proteomic profiling of human atherosclerotic plaques associates IgG with disease severity and outcome

To identify potential factors directly involved in atherosclerosis, we collected atherosclerotic coronary tissue from 98 coronary artery disease (CAD) patients undergoing simultaneous coronary artery bypass graft (CABG) surgery and coronary endarterectomy (CE). Segments were stratified into peripheral (mild plaque) and core (severe plaque) zones, yielding 91 mild and 110 advanced lesions (**Fig. 1A**). For controls, 32 healthy coronary segments were obtained from 14 heart-transplant donors. In total, 233 specimens were subjected to data-independent acquisition (DIA) proteomic profiling (**Fig. 1B** and **Table 1**). Principal component analysis (PCA) revealed a clear distinction among the three groups, validating sampling strategy (**Fig. S1A**). Notably, all four IgG heavy-chain isoforms (IGHG1, IGHG2, IGHG3 and IGHG4) were elevated in atherosclerotic plaques, with the highest in the severe core region (**Fig. 1C**). The broader IgG repertoire, encompassing both light and heavy chains, followed the same pattern (**Fig. S1B**). We next investigated the association of tissue IgG levels with clinical outcomes. Kaplan–Meier analysis revealed elevated plaque levels of IGHG1, IGHG2, and IGHG4 to associate with a trend toward shortened event-free survival, whereas higher IGHG3 was associated with a more favorable course **(Fig. 1D**). Immunohistochemical (IHC) analysis of human coronary arteries corroborated the proteomic data, showing marked IgG deposition within the intima of atherosclerotic lesions (**Fig. 1E, Fig. S2A, B**).

**Figure 1.**
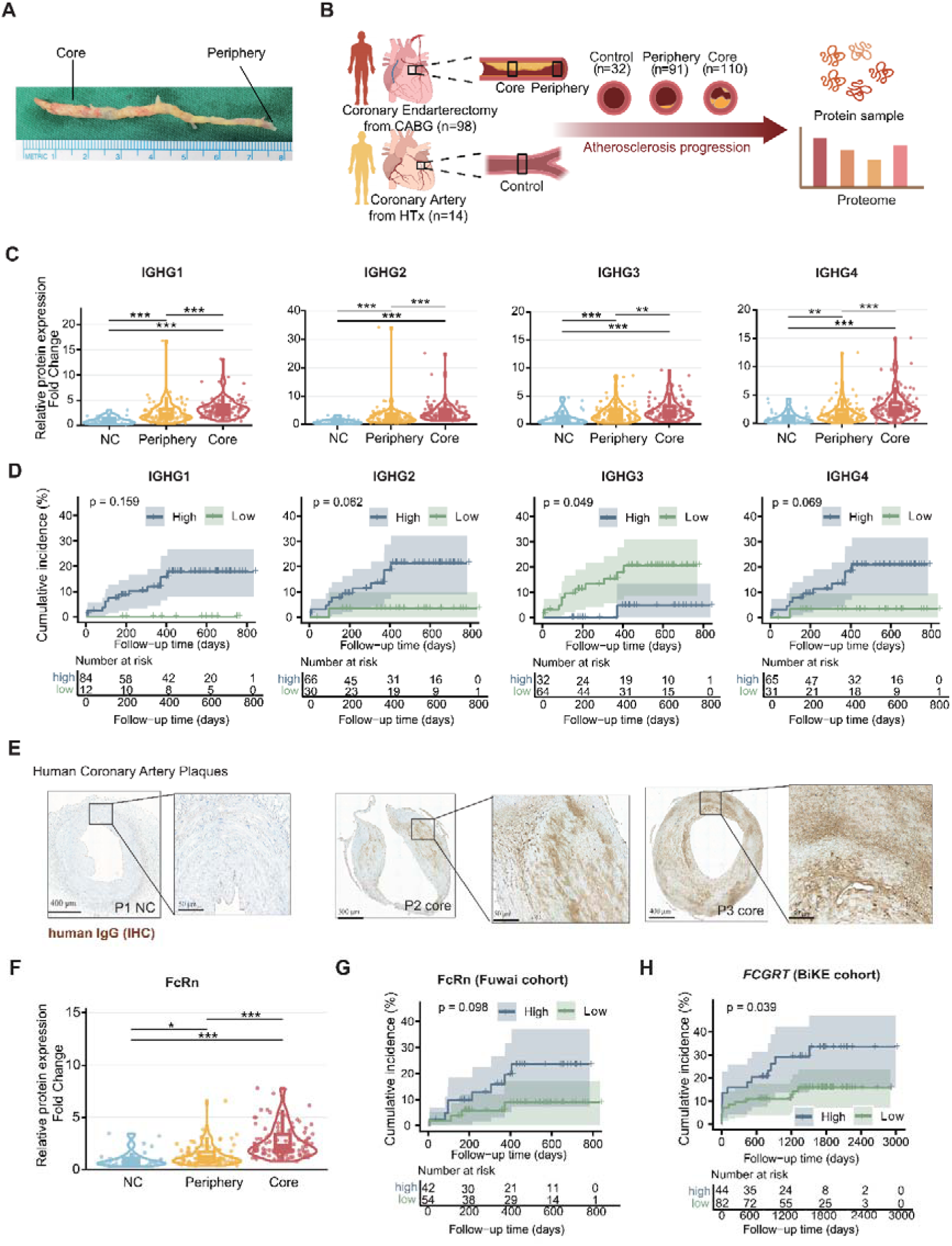
Elevated IgG and FcRn in advanced coronary artery plaques correlate with adverse clinical outcomes. **A**, Representative image of endarterectomized coronary intimal tissue collected during coronary artery bypass graft surgery in patients with coronary artery disease (CAD). **B,** Schematic illustration of the experimental design and cohort stratification. **C,** Violin plots showing the protein abundance of the four IgG heavy chain isoforms (IGHG1–IGHG4) among the three groups. **D,** Kaplan–Meier curves illustrating the association between IgG levels and the incidence of major adverse cardiac and cerebrovascular events (MACCE) over a median follow-up period of 44 months. **E,** Representative immunohistochemical staining of IgG on endarterectomized coronary intimal tissue sections collected during coronary artery bypass grafting (CABG) from patients with coronary artery disease (CAD), (P1, P2, P3 = patient 1, 2 and 3). Scale bars, 50, 300 or 400 μm as indicated. **F,** Violin plots showing FcRn protein expression in plaque cores, peripheral regions, and normal controls. **G-H,** Kaplan–Meier analyses depicting correlations between FCGRT(FcRn) expression and the incidence of MACCE over a median follow-up period of 44 months (**G,** Fuwai cohort) or 3000 days (**H,** BiKE cohort). Data are presented as mean ± SEM. All datasets were assessed for normality and equal variances. Kruskal-Wallis test combined with Dunn’s post-hoc test was used for group comparisons. Survival analyses were performed using Kaplan–Meier method. *p < 0.05, **p < 0.01, ***p < 0.001. See also Figures S1 and S2.

**Table 1.** Baseline characteristics of coronary heart disease (CHD) patients and non-CHD controls.

| Characteristics | CHD (n=98)<sup>a</sup> | Non-CHD (n=14)<sup>a</sup> | P-value<sup>b</sup> |
| --- | --- | --- | --- |
| Age, y | 65 (36-79) | 45 (7-63) | <0.001 |
| Sex, male, n (%) | 67 (68%) | 9 (64%) | 0.8 |
| BMI, y | 25.8 (18.9-37.4) | 21.5 (13.8-27.5) | <0.001 |
| Hypertension, n (%) | 74 (76%) | 3 (21%) | <0.001 |
| Diabetes, n (%) | 45 (46%) | 5 (36%) | 0.5 |

<sup>a</sup>Median (Min-Max); n (%)<sup>b</sup>Wilcoxon rank sum test; Fisher's exact test; Pearson's Chi-squared test**Table 2. Median expression levels of *FCGRT* and IgG in macrophage (Mf) clusters 1-5.**

IgG’s tissue accumulation is dependent on its recycling receptor FcRn and not B cell production^22,24^. Interestingly, FcRn expression mirrored the pattern found with IGHG, displaying stepwise increases in mild and advanced plaques compared to controls (**Fig. 1F**). Transcriptomic interrogation of four independent public data sets, spanning early and advanced lesions, stable and ruptured plaques, and plaques with or without intraplaque hemorrhage (IPH), consistently showed elevated *FCGRT* (encoding FcRn) expression in more severe plaques (**Fig. S2C**). Clinically, low FcRn protein levels were protective against cardiac events in our cohort (**Fig. 1G**). This association was recapitulated in the transcriptomic Biobank of Karolinska Endarterectomy (BiKE) cohort, where higher plaque *FCGRT* correlated with an elevated risk of cerebrovascular events **(Fig. 1H**). Collectively, these cross-cohort human data indicate that plaque IgG, likely facilitated by FcRn as it mediates its recycling, positively correlates with the progression of and adverse outcomes in CAD, highlighting its potential importance in atherosclerosis.

### Single-cell resolution of IgG and FcRn in human atherosclerotic plaques

To explore the cell-specific roles of IgG and FcRn in human plaques, we analyzed a dataset combining cellular indexing of transcriptomes and epitopes by sequencing (CITE-seq) and single-cell RNA sequencing (scRNA-seq) of human carotid atherosclerotic plaques^27^. Plaques were stratified by disease severity (symptomatic vs. asymptomatic; **Fig. 2A**), and cell populations were clustered based on transcriptional profiles (**Fig. 2B**). Interrogating these clusters for *FCGRT* revealed a predominant expression in macrophages, with lower levels in endothelial cells and fibroblasts (**Fig. 2C**). Macrophages in human plaques are heterogenous. Among the five annotated macrophage subsets (Mf1–Mf5; **Fig. S3A**), *FCGRT* expression was highest in clusters 1 and 5 (**Fig. 2D**, and Table 2). Cluster 1, marked by complement component proteins (*C1QA*, *C1QB*, *C1QC*), *CD64*, and *MHCII*, and cluster 5, marked by *C1Q*A-C, immunoglobulin light chain, *MHCII*, and *CD32*, also exhibited the highest surface IgG Fc levels in the CITE-seq data (**Fig. 2E** and **Table 2**), suggesting macrophage FcRn-dependent IgG retention in the plaque microenvironment.

**Figure 2.**
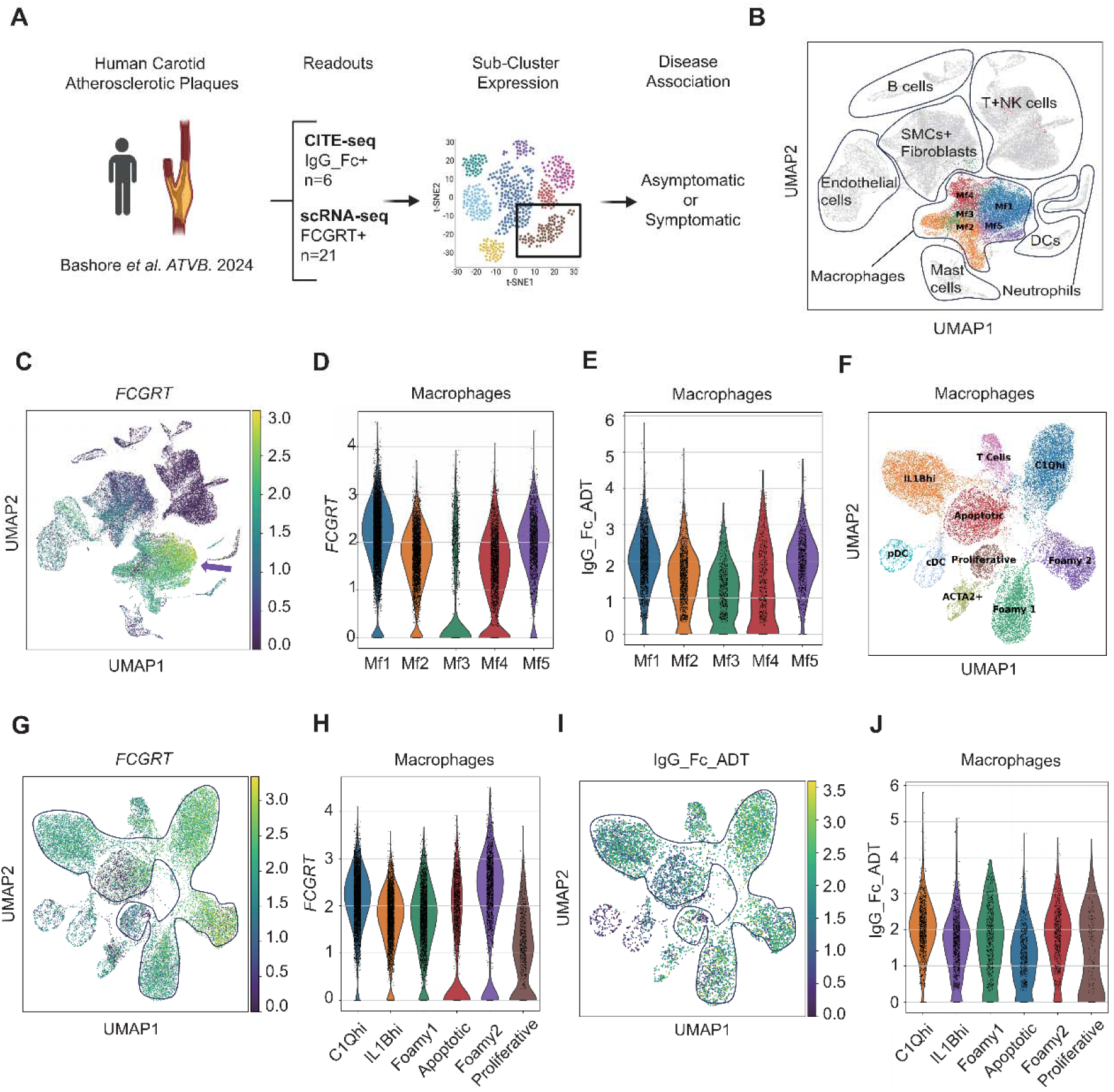
Immune profiling of FcRn and IgG expression in human carotid atherosclerotic plaques. **A**, Schematic overview of the experimental design used to obtain scRNA-seq (n=21) and CITE-seq (n=6) datasets of human carotid atherosclerotic plaques for profiling *FCGRT* and IgG Fc expression. **B,** Uniform manifold approximation and projection (UMAP) of all cell types identified within the collected plaque samples. **C,** UMAP showing *FCGRT* expression levels across all cell types identified within the collected plaque samples. The arrow highlights macrophage populations defined in the referenced study. **D,** Violin plot comparing median *FCGRT* expression across all macrophage clusters. **E,** Violin plot comparing median surface IgG Fc expression across all macrophage subclusters. **F,** UMAP of macrophages and dendritic cells (DCs) clusters colored by their phenotypic and functional characteristics as depicted in the referenced study. pDC, plasmacytoid dendritic cell; cDC, conventional dendritic cell; the T cell cluster represents residual cells. **G,** UMAP of the same clusters indicating the range of *FCGRT* expression from scRNA-seq data. **H,** Violin plots showing median *FCGRT* expression across all macrophage phenotype subclusters. **I,** UMAP of macrophage phenotype clusters (encircled) indicating the range of surface IgG Fc expression from CITE-seq data. **J,** Violin plots showing median IgG Fc expression across all macrophage phenotype subclusters. See also Figures S3.

**Table 2.** Median expression levels of *FCGRT* and IgG in macrophage (Mf) clusters 1-5.

| Feature | Mf1 | Mf2 | Mf3 | Mf4 | Mf5 |
| --- | --- | --- | --- | --- | --- |
| *FCGRT* | 2.18 | 1.73 | 0.00 | 1.45 | 2.03 |
| IgG_Fc_ADT | 2.03 | 1.52 | 1.09 | 1.14 | 2.01 |

While *FCGRT* expression did not differ between symptomatic and asymptomatic plaques (**Fig. S3B–G**), IgG Fc surface levels were elevated in symptomatic patients—specifically 1.76-fold higher in all macrophage subsets when combined (**Fig. S3H**), with the most pronounced increase (1.97-fold) in cluster 5 (**Fig. S3I-M**). To further dissect this heterogeneity, we evaluated *FCGRT* and IgG Fc levels across phenotypic subsets (**Fig. 2F**) defined by distinct gene signatures (**Fig. S3N**). Notably, both *FCGRT* and IgG Fc were most abundant in IL1B^hi^, C1Q^hi^, and foamy macrophages (**Fig. 2G–J** and **Table 3**), suggesting that IgG accumulation is particularly associated with pro-inflammatory and lipid-laden macrophages within the plaque microenvironment.

**Table 3.** Median expression levels of *FCGRT* and IgG Fc in macrophage phenotype clusters.

| Feature | C1Qhi | IL1Bhi | Foamy 1 | Apoptotic | Foamy 2 | Proliferative |
| --- | --- | --- | --- | --- | --- | --- |
| *FCGRT* | 2.19 | 1.79 | 1.77 | 1.28 | 2.31 | 1.02 |
| IgG_Fc_ADT | 2.07 | 1.61 | 1.72 | 1.23 | 1.89 | 1.40 |

### IgG accumulates in atherosclerotic lesions in a macrophage FcRn-dependent manner

To investigate the accumulation of IgG and its role in atherosclerosis, we utilized *Ldlr ^-/-^* mice fed a western diet (WD). Robust IgG deposition was revealed in atherosclerotic lesions (**Fig. 3A, B**), particularly IgG1 and IgG3 subclasses (**Fig. S4A-B**). In contrast, plaque IgM was minimal—likely due to structural constraints of its pentameric form^28^—and IgA deposits were sparse (**Fig. 3A, B**). FcRn expression was apparent and colocalized with IgG-positive regions in the intima (**Fig. 3C** and **Fig. S4C**). Interestingly, exogenous IgG purified from healthy mice preferentially accumulated in plaque-rich regions within the aorta, suggesting plaque IgG is under dynamic exchange towards the arterial wall and unlikely requires autoantibodies (**Fig.3D**). Given that macrophages play a central role in atherogenesis and highly express FcRn, we hypothesized that macrophage FcRn drives plaque IgG accumulation. To test this, we employed myeloid-specific FcRn knockout mice (mKO) and transplanted their bone marrow into irradiated *Ldlr^-/-^* recipients. After 16 weeks of WD feeding (**Fig. 3E**), mKO-transplanted mice exhibited significantly reduced plasma IgG (but not IgM) levels (**Fig. 3F**) and a near-complete loss of IgG in aortic arch (**Fig. 3G, H**) and root lesions (**Fig. 3I, J**). These results recapitulate the accumulation of IgG seen in human plaques and demonstrate its dependence on macrophage FcRn.

**Figure 3.**
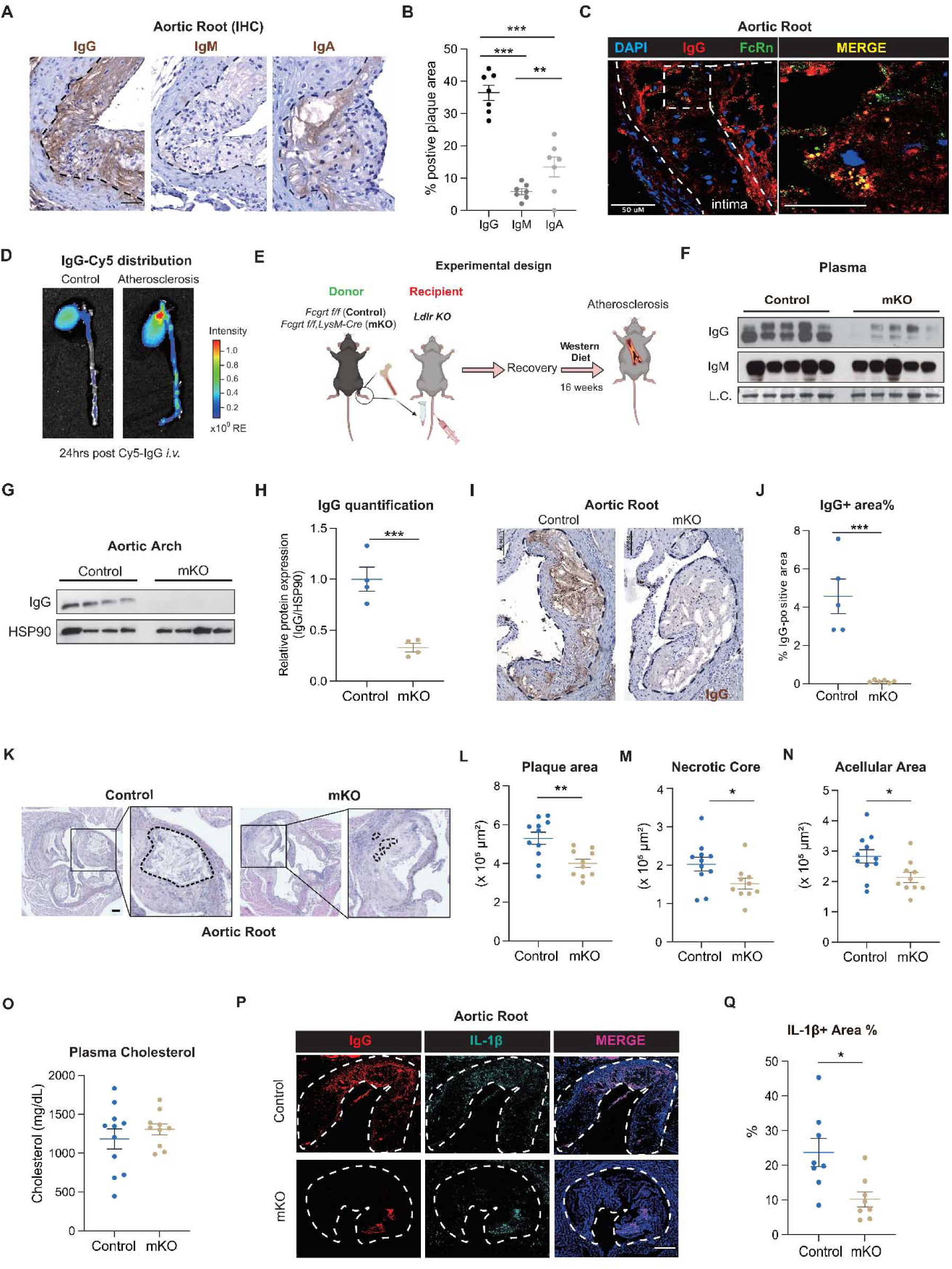
Myeloid FcRn drives plaque IgG accumulation and atherosclerosis progression. **A**, Representative immunohistochemistry (IHC) images showing IgG, IgM, and IgA staining in aortic root lesions from *Ldlr^-/-^* mice fed a western diet (WD) for 16 weeks. Scale bar, 50 μm. Black dashed lines delineate the intima encompassing a plaque-rich leaflet from the medial layer of the aortic root. **B,** Quantification of the percentage of immunoglobulin-positive (Ig+) area within the intimal layer of the aortic root (n=7, 7, 7). **C,** Immunofluorescent staining (IF) for FcRn and IgG in atherosclerotic lesions of *Ldlr^-/-^* mice. Nuclei were stained with DAPI. Scale bars, 50 μm (left) and 25 μm (right). White dashed lines separate the intima (right) encompassing a plaque-rich leaflet from the medial layer (left) of the aortic root. **D,** Ex vivo fluorescence imaging showing mouse IgG-Cy5 distribution in the aorta of atherosclerotic mice and normocholesterolemic controls. **E,** Schematic of the bone marrow transplantation (BMT) atherosclerosis model. Bone marrow cells from FcRn myeloid knockout (mKO) or control mice were transplanted into *Ldlr^-/-^* recipients, which were then fed a WD for 16 weeks to induce atherosclerosis after recovery. **F,** Plasma levels of IgG and IgM in control and mKO mice as determined by western blotting (WB); LC = loading control (Coomassie Brilliant Blue). **G-H,** IgG protein levels in the aortic arch of BMT atherosclerotic mice, as measured by WB (**G**) and quantified in (**H**) (n=4, 4). HSP90 was used as a loading control. **I,** Representative IHC images of IgG staining in the aortic root plaques from BMT mice. Scale bar, 100 μm. **J**, Quantification of the IgG-positive area within the intima, encircled in black (n=5, 5). **K,** Representative H&E staining of aortic root sections from mKO and control BMT *Ldlr^-/-^* mice with atherosclerosis. Scale bar, 100 μm. **L,** Quantification of aortic root lesion area (n=11, 10). **M-N,** Quantification of the necrotic core area (**M**) and acellular area (**N**) within plaques, as delineated by the dashed circle in (**L**) (n=11, 10). **O,** Total plasma cholesterol levels in mKO and control BMT mice with atherosclerosis at sacrifice (n=11, 10). **P**, Immunostaining of IgG and IL-1β in the intima region (encircled in white) of aortic root lesions from BMT mice. Scale bar, 100 μm. DAPI was used for nuclear staining. **Q**, Quantification of total IL-1β+ area out of the total intima region (n=8, 8). Data represent mean ± SEM of independent biological replicates. All datapoints were assessed for normality and equal variances. Statistical analyses were performed using One-way ANOVA in (**B**). Two-tailed Student’s *t*-tests were used for comparisons between two groups. Mann-Whitney *U* test was used for lesion analyses that did not pass normality. **p < 0.01, ***p < 0.001. See also Figures S4-S5.

### Inhibiting plaque IgG accumulation protects against atherosclerosis

To determine whether FcRn-mediated IgG accumulation influences atherosclerosis outcome, we evaluated plaque morphometrics in this cohort. Myeloid-specific deletion of FcRn reduced aortic root lesion area by over 30%, as well as plaque necrosis—a hallmark of unstable, rupture-prone lesions^5^—and total acellular area (**Fig. 3K-N**). These benefits occurred without changes in body weight, glucose tolerance or total and non-HDL cholesterol (**Fig. 3O, Fig. S4D-G**) but along with an increase in circulating HDL (**Fig. S4H-J**).

To validate these findings, we employed a PCSK9-overexpression model (AAV-PCSK9) of hypercholesterolemia in mKO and control mice (**Fig. S5A**). Consistently, mKO mice exhibited reduced plaque and circulating IgG and a trending ∼30% decrease in aortic root lesion area after 16-wk WD feeding (**Fig. S5B-E**). Necrotic and acellular areas were consistently and significantly smaller, again without alterations in cholesterol levels (**Fig. S5F-I**). In the BMT model, mKO-transplanted *Ldlr^-/-^* mice showed normal glucose tolerance and insulin sensitivity due to restrained weight gain after irradiation (**Fig. S4D**). In contrast, AAV-PCSK9-overexpressing mKO mice gained less weight on WD and displayed improved metabolic parameters (**Fig. S5J–L**), recapitulating previous observations in obesity^24^. Crucially, both models demonstrated comparable attenuation of atherosclerosis —irrespective of metabolic differences—underscoring a direct pro-atherogenic role for IgG within the plaque microenvironment.

Next, we sought to evaluate the impact of IgG accumulation on plaque inflammation. In the BMT model, aortic root lesions from mKO-transplanted *Ldlr^-/-^* mice exhibited reduced CD68^+^ macrophage area and fewer Ki67^+^ macrophages (**Fig. S4K-M**), indicating decreased pan-macrophage activation and their subsequent proliferation^29^. Furthermore, the inflammatory cytokine IL-1β was markedly reduced both in plaques and in circulation (**Fig. 4P-Q**). Together, these findings demonstrate that preventing plaque IgG accumulation curbs atherosclerosis burden.

**Figure 4.**
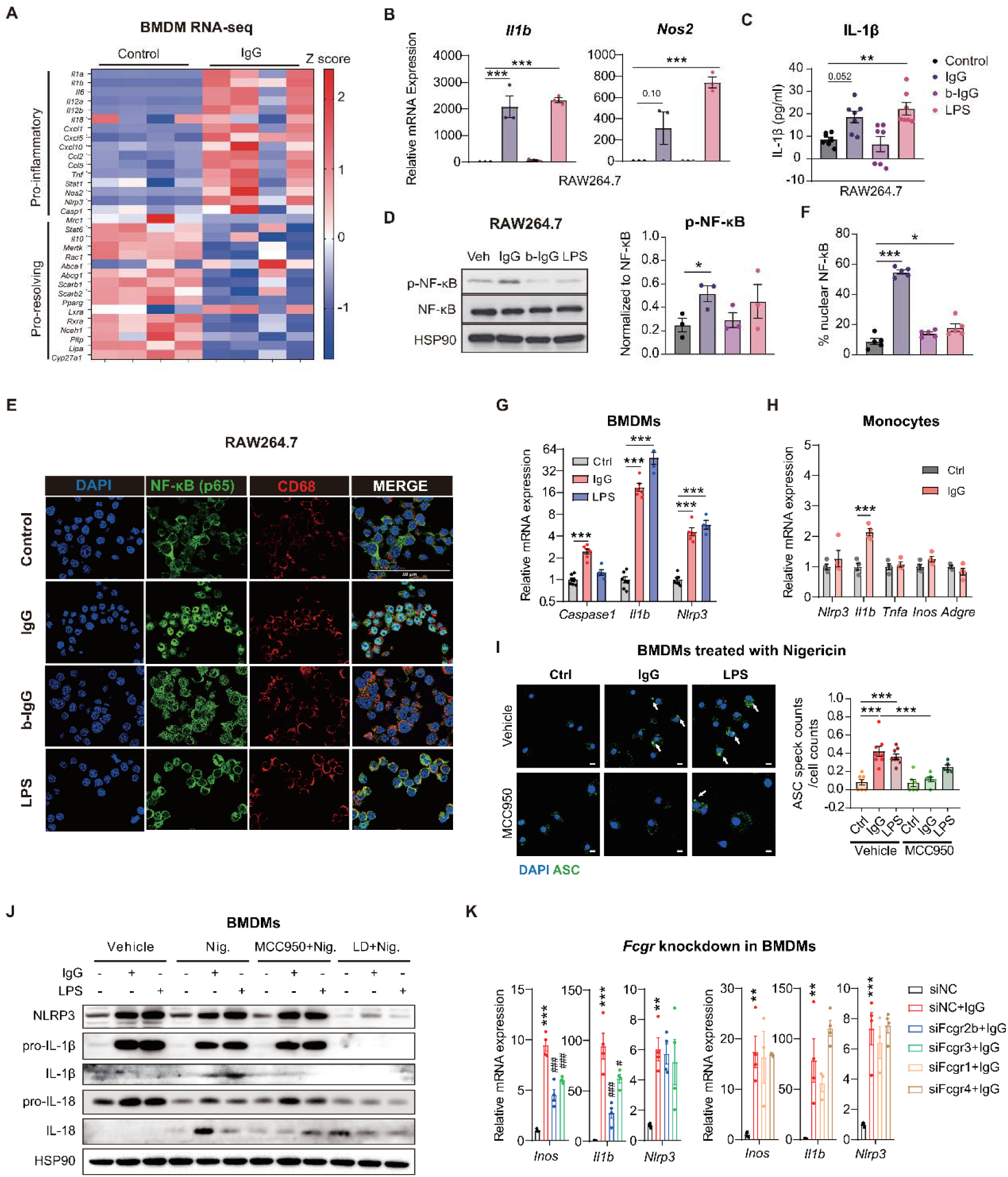
IgG activates NF-κB and the NLRP3 inflammasome in macrophages. **A**, Heatmap showing differentially expressed genes in BMDMs after 16-hr 100 μg/ml IgG treatment (n=4, 4). Gene expression was calculated as log2FPKM and scaled as Z-scores. **B,** QPCR analysis of inflammatory genes *Il1b* and *Nos2* in RAW264.7 cells treated with IgG (100 μg/mL), boiled IgG (b-IgG, 100 μg/mL), or LPS (50 μg/mL) for 24 hours (hrs) (n=3/group). **C,** ELISA quantification of the amount of secreted IL-1β from cells in (**B**) (n=7/group). **D,** WB of phosphorylated and total NF-κB in treated cells as in (**B**) and quantification of the ratio of phosphorylated NF-κB to total NF-κB (n=3/group). HSP90 was used as the loading control. **E-F,** RAW264.7 cells were treated with IgG (100 μg/mL), b-IgG (100 μg/mL), or LPS (50 μg/mL) for 15 minutes. Cells were fixed and stained with NF-κB (p65) and CD68 as shown in (**E**), and Quantification of the percentage of cells with nuclear NF-κB localization based on images from (**F**) (n=5/group). Scale bar, 50 μm. **G,** QPCR analysis of *Caspase1, Il1b* and *Nlrp3* in BMDMs treated with IgG (100 μg/mL), or LPS (50 μg/mL) for 24 hrs (n=5/group). **H,** QPCR analysis of *Nlrp3, Il1b, Tnfa, Inos* and macrophage marker *Adgre1*(F4/80) in bone marrow-derived monocytes treated with IgG (100 μg/mL) for 24 hrs (n=5/group). **I,** Representative IF images of BMDMs treated with IgG (100 μg/mL), LPS (50 μg/mL) and MCC950 (100 nM) for 24 hrs, followed by incubation with Nigericin (10 μM) for 30 minutes, and stained for ASC. Scale bar, 20 μm, and the quantification of the number of ASC specks per cell (n=8/group). DAPI was used for nuclear staining and cell counting. **J,** Representative WB analysis of BMDMs treated with IgG (100 μg/mL), LPS (50 μg/mL), MCC950 (100 nM), or Licochalcone D (10 μM) for 24 hrs, followed by Nigericin (10 μM) for 30 minutes. Blots were probed for NLRP3, IL-1β, and IL-18. **K,** QPCR analysis of *Nos2, Il1b*, and *Nlrp3* mRNA levels in WT BMDMs transfected with siRNAs against mouse *Fcgr1, Fcgr2b, Fcgr3, Fcgr4* or sham control(siNC). These cells were treated with IgG (100 μg/mL) for 24 hrs (n=4/group). *Compared with siNC group, #compared with siNC+IgG group. Data represent mean ± SEM. All datasets were assessed for normality and equal variances. One-way ANOVA with multiple comparisons and two-tailed Student’s t-tests for pairwise group comparisons were used for statistical analysis. *^/#^p < 0.05, **^/##^p < 0.01, ***^/###^p < 0.001. See also Figures S6.

### IgG induces an inflammatory milieu in macrophages

Given the marked changes in plaque macrophage burden, we investigated whether IgG directly impacts their inflammatory response ex vivo. Bulk RNA-seq of IgG-treated bone marrow-derived macrophages (BMDMs) revealed a pronounced pro-inflammatory shift, alongside a suppression of genes involved in cholesterol efflux, efferocytosis, and lysosomal activity (**Fig. 4A**). This inflammatory phenotype was replicated in RAW264.7 macrophages, where IgG induced *Nos2* and *Il1b* expression to levels comparable to Lipopolysaccharide (LPS) stimulation (**Fig. 4B**). Notably, boiled IgG (b-IgG) failed to activate these genes. Further, IgG—but not denatured IgG—increased IL-1β secretion (**Fig. 4C**), confirming that native IgG is required for the pro-inflammatory induction of macrophages.

NF-κB governs macrophage inflammation by phosphorylation-dependent nuclear translocation^30,31^, with implications in atherosclerosis^32^. IgG treatment in RAW264.7 cells induced phosphorylation of NF-κB (p65) at Ser536 (**Fig. 4D**) and triggered its rapid nuclear translocation within 15 minutes—a response delayed with LPS and absent with denatured IgG (**Fig. 4E, F** and **Fig. S6A**). NLRP3 inflammasome activation occurs downstream of NF-κB and contributes to the pathogenesis of atherosclerosis^33^. IgG administration induced an upregulation of *Nlrp3*, *Il1b*, and *Caspase1* transcripts, similar to LPS (**Fig. 4G**). Of note, monocytes exhibited only a minimal response to IgG stimulation (**Fig. 4H**), indicating that primed and differentiated macrophages are required for responding to IgG, mirroring the plaque microenvironment. As an indicator of inflammasome activation^34^, ASC speck formation was enhanced by IgG treatment in BMDMs with LPS as a positive control, while this effect was blocked by the NLRP3 inhibitor MCC950 (**Fig. 4I**). Consistently, intracellular levels of NLRP3, pro-IL-1β, and pro-IL-18 were induced by IgG (**Fig. 4J**). Nigericin (signal 2) facilitated the proteolytic cleavage of these pro-forms into mature forms (IL-1β and IL-18)^35^, and this process was effectively suppressed by pharmacological inhibition of NLRP3 by MCC950 or NF-κB by Licochalcone D (LD) (**Fig. 4J**). Our results show that IgG can effectively prime and activate the NLRP3 inflammasome.

To understand regulatory pathways, we pretreated BMDMs with various kinase inhibitors (MEK: PD98059; PI3K: LY294002; Src/BCR-ABL: Dasatinib; BTK: Ibrutinib; Syk: R406) prior to IgG exposure. None abolished IgG’s induction of inflammasome-associated genes, with some even exacerbating the response (**Fig. S6B**). Furthermore, siRNA-mediated knockdown of FcγRs (*Fcgr1*, *Fcgr2b*, *Fcgr3*, and *Fcgr4*) in BMDMs only partially or minimally attenuated pro-inflammatory gene induction (**Fig. 4K, Fig. S6C**). Together, these findings demonstrate that IgG stimulates a pro-inflammatory state in macrophages primarily through NF-κB, eliciting downstream activation of the NLRP3 inflammasome. A similar trend was observed in THP-1 induced human macrophages (**Fig. S8H, I**).

### IgG directly activates TLR4 in macrophages independently of its antigen-neutralization function

TLR4 is a prominent upstream activator of NF-κB signaling^36,37^. In RAW264.7 macrophages, IgG-induced NF-κB phosphorylation and nuclear translocation were completely blocked by the TLR4 inhibitor TAK242, (**Fig. 5A, B**; and **Fig. S6D, E**), as was the induction *of Nos2* and *Il1b* (**Fig. 5C**), IL-1β secretion (**Fig. S6F**), and NF-κB phosphorylation (**Fig. 5D**). Similar dependences on TLR4, NF-κB, and NLRP3 were observed in BMDMs and human THP-1 macrophages (**Fig. S6G-I**). Interestingly, pro-IL-1β and IL-1β levels were similarly induced by a monoclonal IgG antibody against PD1 (hPD1 mAb), and this effect was also blocked by TAK242 (**Fig. 5E**), implying that antigen recognition is not required for TLR4 activation. To test this directly, we treated BMDMs with IgG’s antigen-binding fragment Fab or constant Fc fragment. Phosphorylation and nuclear translocation of NF-κB were largely recapitulated by Fc but not Fab treatment (**Fig. 5F; Fig. S7A, B**) accompanied by upregulation of downstream targets (NLRP3, pro-IL-1β, and pro-IL-18) (**Fig. 5G; Fig. S7C**). Fc-induced activation was also abolished by TAK242. Moreover, treatment of mice with an IgG Fc fragment was sufficient to enrich its presence in atherosclerotic plaques (**Fig.5H**). Overall, we show that IgG plays a pathogenic role in the activation of macrophages and the progression of atherosclerosis in a noncanonical manner.

**Figure 5.**
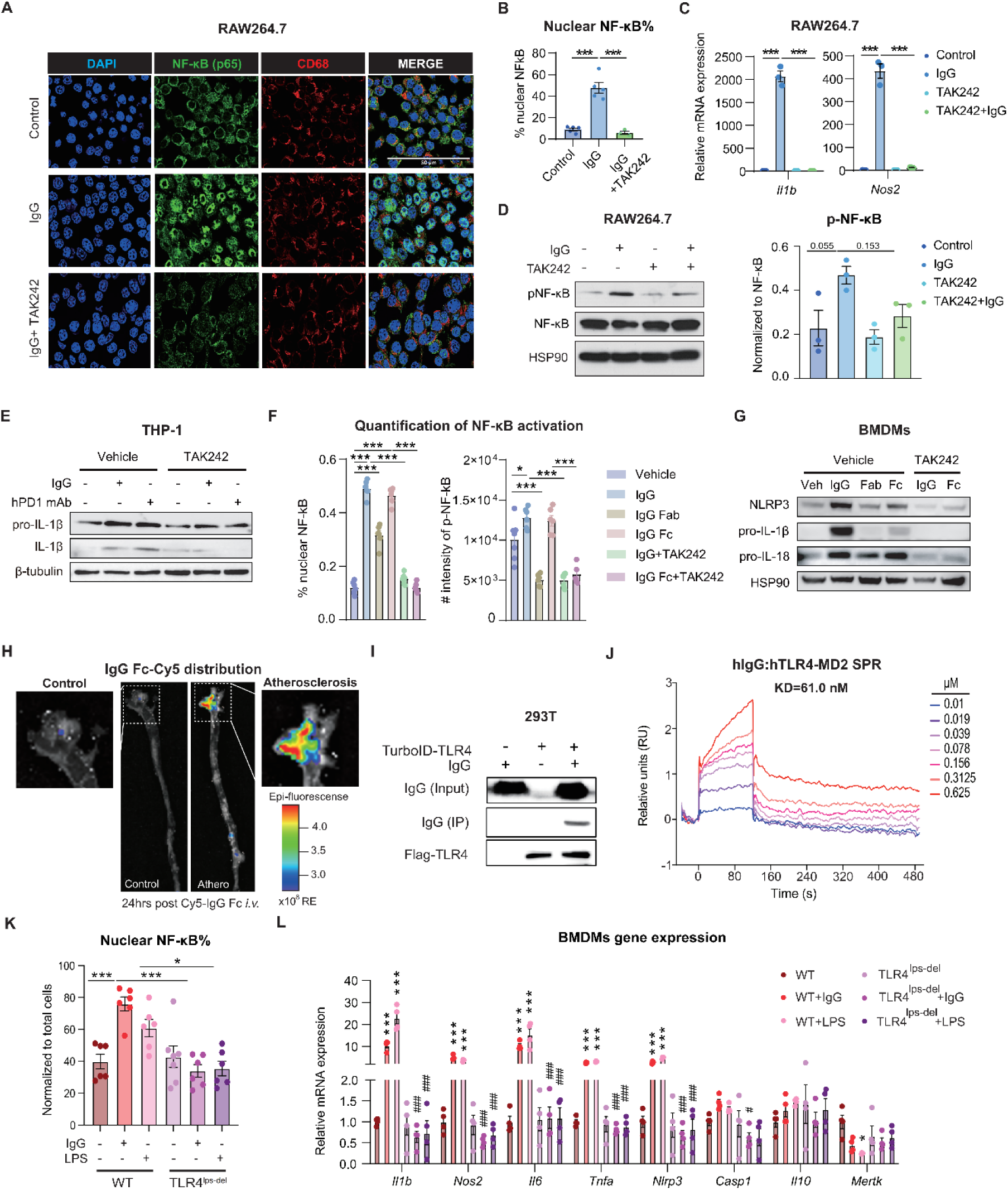
IgG stimulates TLR4 signaling upstream of NF-κB. **A**, Representative IF images of NF-κB (p65) and CD68 in RAW264.7 cells pretreated with TLR4 inhibitor TAK242 (1 μM), followed by IgG (100 μg/mL) stimulation for 2 hrs. Scale bar, 50μm. **B,** Quantification of percentage of nuclear NF-kB cells based on images from (**A**) (n=5, 5, 3). **C,** QPCR analysis of *Il1b* and *Nos2* expression in cells treated as in (A) for 24 hrs (n=3/group). **D,** WB analysis of phosphorylated and total NF-κB in RAW264.7 cells treated as in (**C**) and quantification of phosphorylated NF-κB levels (n=3/group). HSP90 was used as the loading control. **E**, WB analysis of pro-IL-1β, IL-1β in lysates from human THP-1 cells pretreated with TAK242 (1 μM) for 1 hr, followed by 12-hour treatment with vehicle, IgG (100 μg/mL) or hPD1 mAb (100 μg/mL). **F**, Quantification of phospho-NF-κB intensity and the NF-κB p65 nuclear translocation based on IF images of BMDMs pretreated with TAK242 (1 μM) for 1 hr, followed by 1-hr treatment with vehicle, IgG (100 μg/mL), IgG Fab domain (20 μg/mL), IgG Fc domain (100 μg/mL) as shown in Extended Data Figure 9A and 9B (n=8/group). **G**, WB analysis of NLRP3, pro-IL-1β, pro-IL-18 in lysates from BMDMs pretreated with TAK242 (1 μM) for 1 hr, followed by 1-hr treatment with vehicle, IgG (100 μg/mL), IgG Fab domain (20 μg/mL), IgG Fc domain (100 μg/mL). **H,** Ex vivo fluorescence imaging showing mouse IgG Fc-Cy5 distribution in the aorta of control and atherogenic mice. **I,** WB analysis of streptavidin pull-downs from TurboID-TLR4/MD2 co-transfected 293T cells. Cells were treated with vehicle or 100 μg/mL IgG for 12 hrs, 1 day post transfection. **J,** Surface plasmon resonance analysis of kinetic binding of human IgG (hIgG) with human TLR4-MD2 complex. The titration of native IgG at concentrations from 10nM – 625nM. **K,** Quantification of percentage of cells with nuclear NF-kB localization from IF images shown in (**J**) (n=6, 6, 7, 6). **L,** QPCR analysis of inflammatory gene expression in WT and TLR4^lps-del^ BMDMs (n=4/group). *Compared with WT group, ^#^ compared with TLR4^lps-del^ group. Data represent mean ± SEM. All datasets were assessed for normality and equal variances. One-way ANOVA followed multiple comparisons was used for statistical analysis. *^/#^p < 0.05, **^/##^p < 0.01, ***^/###^p < 0.001. See also Figures S7-S8.

**Figure 6.**
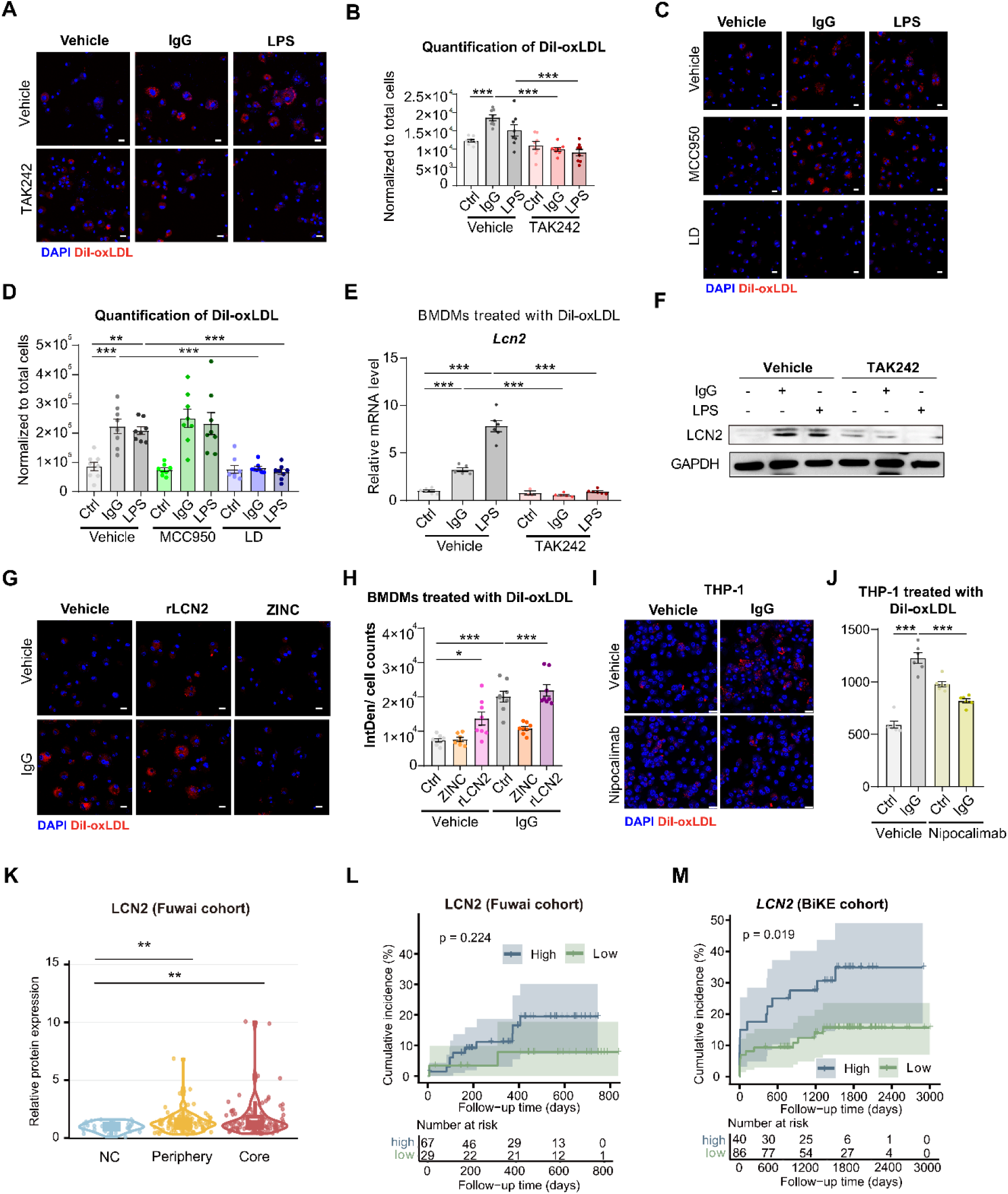
IgG promotes macrophage foam cell formation through a TLR4/NF-κB/LCN2 axis. **A**, Representative IF images of Dil-oxLDL staining in BMDMs treated with Vehicle, IgG (100 μg/mL), LPS (50 μg/mL) in the presence or absence of TAK242 (1 μM) for 18 hrs, followed by incubation with Dil-oxLDL (10 μg/mL) for 6 hrs. Cells were fixed and stained for imaging. Scale bar, 20 μm. **B,** Quantification of Dil-oxLDL fluorescence intensity per cell from images in (**A**) (n = 8 per group). **C,** Representative IF images of Dil-oxLDL in BMDMs treated with IgG (100 μg/mL), LPS (50 μg/mL), MCC950 (100 nM), or Licochalcone D (10 μM) for 18 hrs, followed by Dil-oxLDL (10 μg/mL) for 6 hrs. scale bar, 20 μm. **D,** Quantification of Dil-oxLDL fluorescence intensity per cell from (**C**) (n = 8 per group). **E,** QPCR analysis of *Lcn2* expression in BMDMs treated with oxLDL (50 μg/mL), followed by IgG (100 μg/mL), LPS (50 μg/mL) or TAK242 (1 μM) for 24 hrs (n=5, 5, 5, 3, 5, 5). **F,** Representative WB analysis of BMDMs from (**E**). LCN2 was probed. **G,** Representative IF images of Dil-oxLDL in BMDMs treated with IgG (100 μg/mL), the LCN2 inhibitor ZINC00640089 (1 μM), or recombinant LCN2 (10 μg/mL) for 18 hrs, followed by Dil-oxLDL (10 μg/mL) for 6 hrs. Scale bar, 20 μm. **H,** Quantification of Dil-oxLDL fluorescence intensity per cell from images shown in (**G**) (n = 8 per group). **I,** Representative IF images of Dil-oxLDL in THP-1 cells treated with Vehicle, IgG (200 μg/mL), in the presence or absence of Nipocalimab (10 μg/mL) for 18 hrs, followed by Dil-oxLDL (10 μg/mL) for 6 hrs. **J,** Quantification of Dil-oxLDL fluorescence intensity per cell from (**I**) (n = 6 per group). **K,** Proteomic analysis of LCN2 protein levels in normal arteries, plaque periphery and core regions from the Fuwai cohort. **L,** Kaplan–Meier curves showing the association between LCN2 expression and the incidence of MACCE over a median follow-up period of 44 months from the Fuwai cohort. **M,** Kaplan–Meier curves depicting the association between *LCN2* and the incidence of MACCE over a median follow-up period of 3000 days from the BiKE cohort. Data represent mean ± SEM. All datasets were assessed for normality and equal variances. Statistical comparisons were performed using One-way ANOVA with multiple comparisons. *p < 0.05, **p < 0.01, ***p < 0.001.

To determine whether IgG directly engages with TLR4, we employed TurboID proximity labeling in 293T cells, revealing positive labelling of IgG by TurboID-tagged TLR4 (**Fig. 5I**). Surface plasmon resonance (SPR) confirmed a robust binding between human IgG (hIgG) and the hTLR4-MD2 co-receptor complex (**Fig. 5J; Fig. S8A**). In contrast, IgG failed to bind to TLR4 without MD2 (**Fig. S8B, C**), highlighting that an integral TLR4 complex is required. In BMDMs derived from TLR4^lps-del^ mice, which are deficient in TLR4 signaling^38^, IgG failed to induce NF-κB’s nuclear translocation (**Fig. 5K, Fig. S8D**) or phosphorylation (**Fig. S8E-G**). Similarly, IgG’s induction of pro-inflammatory genes (*Il1b*, *Nos2*, *Il6*, *Nlrp3*) and repression of pro-resolving genes (*Mertk*) were abolished (**Fig, 5L**), and IL-1β secretion was unchanged in TLR4^lps-del^ BMDMs (**Fig. S8H**). Collectively, these results establish TLR4 as a necessary mediator for IgG to activate NF-κB-driven inflammatory signaling in macrophages.

### IgG promotes macrophage foam cell formation through the TLR4/NF-κB/LCN2 axis

Lipid-laden macrophages, or foam cells, are key determinants of atherosclerosis progression^39^, and their formation is promoted by TLR4 activation^40–42^. We therefore tested whether IgG can activate TLR4 to accelerate this process. BMDMs were primed with IgG treatment and subsequently loaded with Dil-oxLDL. IgG significantly enhanced oxLDL uptake in BMDMs, similar to LPS, and TLR4 inhibition effectively abolished this effect (**Fig, 6A,B**). This IgG-induced foam cell formation was also blocked by the NF-κB inhibitor Licochalcone D but not by the NLRP3 inhibitor MCC950 (**Fig, 6C,D**), indicating its dependence on TLR4/NF-κB signaling but not the NLRP3 inflammasome.

The mediators underlying the transformation of inflammatory macrophages into foam cells are under active investigation. Lipocalin-2 (LCN2) is a secretory protein implicated in immune responses^43,44^ and atherosclerosis^45–47^. Noting its upregulation by IgG treatment in BMDMs by RNA-seq, we validated that it was robustly induced by IgG in a TLR4-dependent manner (**Fig, 6E,F**). Recombinant LCN2 protein facilitated Dil-oxLDL uptake in BMDMs, with no additive effect from IgG, and this process was abrogated by the LCN2 inhibitor ZINC (**Fig, 6G,H**), suggesting that IgG acts through LCN2 to promote macrophage foam cell formation. Interestingly, IgG-induced foam cell formation was also recapitulated in human THP-1 macrophages and blocked by Nipocalimab, a clinically used monoclonal antibody against FcRn (**Fig, 6I, J**). We further investigated the clinical relevance of this axis. In the proteomic Fuwai cohort, LCN2 protein levels were increased in both mild (periphery) and advanced (core) coronary artery plaques compared to controls (**Fig, 6K**), with higher plaque LCN2 trending toward shorter event-free survival (P=0.224) (**Fig, 6L**). In the larger BiKE carotid plaque transcriptomic cohort, high plaque *LCN2* expression was associated with significantly shorter event-free survival (P=0.0194) (**Fig, 6M**). In conclusion, we identify LCN2 as a downstream effector of IgG-induced TLR4/NF-κB signaling that promotes macrophage foam cell formation, suggesting that the robust changes in plaque burden with IgG accumulation inhibition may be occurring through this milieu.

## DISCUSSION

The vascular endothelium is perpetually bathed in abundant immunoglobulins—so ubiquitous that their pathogenic potential is often overlooked. Here, we identify IgG as a critical pro-atherogenic factor, one that accumulates in plaques under dyslipidemia conditions—a finding with direct translational relevance as it correlates positively with atherosclerosis burden in humans. This accumulation is governed by macrophage FcRn-mediated IgG recycling, which not only sustains intraplaque IgG levels but also amplifies its inflammatory impact via TLR4/NF-κB signaling. Inhibiting this accumulation reduces plaque inflammation and attenuates atherosclerosis in mice. Our findings redefine atherosclerosis pathogenesis through the lens of an abundant yet underappreciated mediator, positioning IgG as a novel and potential therapeutic node.

While IgG is indispensable for immune defense, its unchecked accumulation in plaques—akin to its recently described role in thrombosis via platelet complement activation^48^—can exacerbate cardiovascular pathophysiology. Notably, pathogenic IgG may arise not only from increased abundance but also from altered specificity or post-translational modifications (e.g., glycosylation^49^), potentially converting IgG from a protective agent to a driver of maladaptive inflammation. Further, different IgG subtypes exert divergent effects. We observed that elevated plaque levels of IgG1, IgG2 and IgG4 were positively correlated with CAD-related events, whereas IgG3 was associated with a more favorable outcome. Intriguingly, IgG3 is the least abundant IgG subtype and has the shortest half-life due to its weak binding affinity to FcRn among all IgG isotypes^50^. Although oxLDL-specific IgG autoantibodies have been documented in plaques^15,51^, regular IgG isolated from healthy mice, which unlikely contain such atherogenic autoantibodies, was actively enriched in plaques, and IgG-Fc, even without IgG’s antigen-recognizing Fab domain, was also able to enter the plaque, confirming that plaque IgG deposition is not necessarily antigen-specific.

The canonical paradigm holds that IgG binds antigen via its Fab to form ICs, which then trigger FcγR-mediated responses^52^. Our results demonstrate that IgG can bypass this classic pathway. IgG directly interacts with TLR4, and its constant Fc domain is sufficient to activate NF-κB in macrophages, indicating antigen-binding is not necessary. This pathway does require the native IgG structure and an intact TLR4/MD2 complex, as neither denatured IgG nor TLR4 alone could replicate this effect. TLR4 is a sentinel of innate immunity, known for its response to bacterial LPS but also to endogenous ligands like S100A8^53^, heat shock proteins (HSPs), LDL and viral proteins^54,55^. Herein, we reveal a non-canonical function of IgG as a direct activator of innate immunity, forging a molecular bridge between the adaptive and innate immune systems. This mechanism provides a streamlined explanation for chronic inflammation in atherosclerosis, with LCN2 identified as a critical downstream mediator promoting plaque foam cell formation. Collectively, our study establishes a new paradigm in which IgG, acting as a pathogenic factor driving atherosclerosis through a non-canonical mechanism that may also apply to other chronic inflammatory diseases.

Systemic IgG homeostasis relies more on FcRn-mediated recycling than *de novo* B cell production^56^. This study pioneers the exploration of IgG recycling in atherosclerosis. By establishing that plaque IgG deposition is not antigen-binding-dependent but rather FcRn-dependent, we unveil FcRn as a previously unrecognized arbiter of disease progression. Clinically, our proteomic analysis links both IgG and FcRn to atherosclerosis severity, a connection also confirmed in mice, where macrophage-specific FcRn deletion reduces plaque IgG depots and attenuates disease outcome. We further endow a dual role for IgG in driving pro-inflammatory macrophage responses and promoting foam cell formation. Our previous studies reveal that mKO mice prevent IgG from accumulating in the serum and in visceral adipose tissue in aging and obesity, but not at basal conditions^57,58^. Therefore, strategically targeting FcRn offers distinct advantages over B-cell depletion, which compromises all antibody production^59,60^, as it selectively reduces IgG while preserving immune competence. Indeed, we show that the FDA-approved FcRn antibody—Nipocalimab works effectively to suppress foam cell formation in activated human THP-1 macrophages. By identifying IgG as a key proatherogenic factor, our work unveils a novel therapeutic avenue to curb cardiovascular risk, particularly in high-risk conditions like aging and obesity.

## Author contributions

T.Z., K.Z., L.W. and L.Q. conceptualized the study, designed the experiments, and wrote the manuscript. T.Z., K.Z., S.H., L.Y., Z.Y., D. L., Q.W., and QF.W. performed the experiments. T.Z., K.Z., S.H., C.X., B.L., X.L., M.S., and A.R.K. performed data analyses. S.H., Z.H. and W.F. were in charge of human sample collection and analyses; F.Y. and M.P.R. helped with resources and reagents. L.Q. is the primary overseer of this study, and as such, has full access to all the data in the manuscript and takes responsibility for the integrity of the data and the accuracy of the data analysis.

## Data, code, and materials availability

- Data that support the findings of this study are available from the corresponding author upon reasonable request.
- Clinical proteomic data will be made publicly available upon publication.
- Mouse RNA-seq data can be found in the GSA with accession number PRJCA020637^22^.
- Human scRNA-seq and CITE-seq data can be found in the Gene Expression Omnibus with persistent ID GSE253904^27^.
- This paper does not contain original code.
- The research materials in this study are available upon request from the lead contact.

## Supplementary Materials

Experimental model and study participant details.

Figures. S1 to S8

Graphic abstract.

