## Supplemental Material for "IgG Propels Atherosclerosis by Noncanonically Activating Macrophages"

**EXPERIMENTAL MODEL AND STUDY PARTICIPANT DETAILS**

### **Methods**

**Sample preparation, LC-MS/MS analysis, and data processing for clinical proteomic profiling**

Tissue pulverization was done in liquid nitrogen, protein extraction was done using a sodium dodecyl sulfate (SDS) Tris‑based buffer for reduction/alkylation, followed by acetone precipitation and purification via ultrafiltration. Protein quantification was performed using the Bradford assay, and quality was confirmed by SDS-PAGE. For LC-MS/MS analysis in data-independent acquisition (DIA) mode, digested peptides were separated using nano-UHPLC and analyzed on an Orbitrap Astral mass spectrometer. Database searching was conducted with DIA-NN library search software (version 1.8.1) against the UniProt human database, and identified proteins were filtered based on a 1% false discovery rate (FDR). Normalization was performed by median centering, and missing values were imputed using k-nearest neighbor (k-NN) after excluding proteins with more than 25% missing data.

**Differential expression analysis in CE cohorts and transcriptomic datasets**

Differential expression analysis was performed on both proteomic data from the Fuwai cohort and transcriptomic data targeting *FCGRT* (FcRn). For the proteomic analysis, samples were categorized into Core, Periphery, and NC groups. In the transcriptomic analysis, samples were divided into two groups based on dataset annotations. In both analyses, expression values were normalized to the mean of a reference group (NC for proteomics) to calculate relative expression or fold change. The Kruskal-Wallis test with Dunn's post-hoc test was used for multiple group comparisons, while the Student's t-test was applied to compare between two groups, with statistical significance determined using an FDR threshold of 0.05.

**Description of Fuwai cohorts and BiKE cohorts with clinical follow-up**

Two cohorts were utilized for this correlation analysis. The first cohort integrated our in-house proteomic data of human coronary arteries with the follow-up cohort from Fuwai Hospital. For the Fuwai Hospital cohort, the follow-up information referred to: the primary outcome was the occurrence of major adverse cardiovascular and cerebrovascular events (MACCE), which was defined as a composite endpoint including all-cause death, nonfatal myocardial infarction, repeat revascularization, and stroke (encompassing both hemorrhagic and ischemic stroke). The second cohort was based on the public dataset GSE21545, which was retrieved from the Biobank of Karolinska Endarterectomy (BiKE) database. The BiKE database contains transcriptional profiles of 126 human carotid plaque samples, which were collected from patients who underwent carotid endarterectomy. Over a median follow-up period of 44 months, 25 MACCE events were documented: 18 ischemic strokes and 7 myocardial infarctions.

#### **Immunohistochemical staining of human coronary arteries**

Atherosclerotic plaque specimens were obtained from coronary artery endarterectomies performed during coronary artery bypass graft (CABG) procedures at Fuwai Hospital, China. Formalin-fixed paraffin-embedding samples containing intimal tissues were processed for histological evaluation. Plaque morphology was assessed by H&E staining, with IgG deposition visualized by immunohistochemistry using a rabbit anti-human IgG antibody (ZSGB-BIO #ZA-0448). Complete patient demographics, clinical characteristics, and cardiovascular imaging findings are presented in Extended Data Figure 1.

#### **scRNA-seq and CITE-seq analyses of human carotid artery plaques**

We analyzed scRNA-seq and CITE-seq data from human carotid atherosclerotic plaques obtained from a published study^27^, which included 21 patients (8 asymptomatic, 13 symptomatic with prior stroke/TIA). CITE-seq data with antibody-derived tags was available for 6 subjects. All data use was approved by Columbia University's IRB (AAAR6796). The original clustering analysis was reproduced, maintaining all identified cell populations (including dendritic cells and T cells) to evaluate *FCGRT* and IgG Fc expression. As noted in the source study, the ACTA2+ cluster comprises both smooth muscle cells and macrophages, while T cells represent a residual population rather than plaque macrophages. These cluster designations were preserved in our analysis to maintain consistency with the original dataset. A pseudobulk approach was employed to compare RNA and protein expression between symptomatic and asymptomatic subjects. For each cell cluster of interest, scRNA-seq read counts and CITE-seq protein counts were aggregated across subjects within each clinical group. Differential expression analysis was performed using DESeq2^61^, retaining features with ≥10 counts across subjects. Significant differences were defined as fold change ≥1.5 (symptomatic/asymptomatic ratio of median normalized counts) with Benjamin-Hochberg adjusted p-value <0.05. Both datasets are available in GEO (GSE253904).

#### **Metabolic studies in mice**

For glucose tolerance tests (GTT), 16-hr overnight-fasted mice were injected intraperitoneally with glucose (2 g/kg body weight). Blood glucose levels were measured at 0, 15, 30, 60, 90, and 120 minutes post-injection using a One Touch Ultra glucometer. For insulin tolerance tests (ITT), 4-hour-fasted mice received intraperitoneal insulin (0.7 U/kg body weight), with glucose measurements taken at 0, 15, 30, 45, and 60 minutes. All fasting procedures were conducted in cages with fresh bedding.

**Blood collection and analysis**

Blood was collected via cardiac puncture from euthanized mice and divided into two aliquots. One aliquot was immediately analyzed for complete blood counts, including total white blood cells (WBC) and monocyte populations, using a Cell Blood Counter (Oxford Biosciences). The second aliquot was centrifuged for plasma isolation and subsequent analyses. Plasma lipid profiling included: total cholesterol (Fujifilm Wako), HDL cholesterol (Fujifilm Wako), and calculated non-HDL cholesterol (total cholesterol - HDL cholesterol).

#### **Plasma lipoprotein fractionation**

Pooled plasma samples (n=3-4 mice per round; 3 independent rounds) from BMT mice (mKO and Control) were diluted 1:1, filtered, and subjected to fast-protein liquid chromatography (FPLC) as previously described^62^. Samples were fractionated using a Superdex 200 Increase 10/300GL column (GE Healthcare) with FPLC buffer (100 mmol/L Tris, 0.4 g/L NaN3, pH 7.5) at a flow rate of 0.3 mL/min. Eluted fractions were analyzed for lipoprotein distribution using Total Cholesterol E assay (Fujifilm Wako).

#### **Morphometric analysis of mouse aortic root lesions**

#### Mice were euthanized by CO_2_ asphyxiation and perfused with saline. Aortas were dissected using butterfly scissors, with careful removal of perivascular fat and lymph nodes. Aortic arches were snap-frozen for protein/RNA analysis, while aortic roots were fixed in 10% formalin overnight, transferred to 70% ethanol, and paraffin-embedded. Serial 6 µm sections were mounted on charged slides for histological assessment. Plaque morphology was evaluated in H&E-stained sections (6 slides per sample, 60 µm apart) using a Keyence microscope. Total plaque area, necrotic core size, and acellular regions were quantified across all three valve leaflets as previously described^62^.

**Ex vivo fluorescence imaging detection of IgG and IgG Fc fragment in mouse aortas**

IgG or IgG Fc fragment was conjugated with Cy5 using Lumiprobe NHS-Cy5 (Bidepharm, Cat. No. 1032678-42-4) according to the manufacturer’s instructions. Mice were administered these labeled compounds via intravenous injection (*i.v.*) at a dose of 400 μg per mouse. After sacrifice at 24 hours post-administration, the aorta was isolated and the Cy5 fluorescence signal was quantified using an IVIS Spectrum (PerkinElmer).

**Immunohistochemistry**

Tissue sections (6 µm) from paraffin- or OCT-embedded samples were prepared on charged slides. Paraffin sections were deparaffinized in xylene and rehydrated through an ethanol series (100%→95%→70%). Frozen sections were air-dried, fixed in 4% paraformaldehyde (PFA) (10 min), and PBS-washed. For antigen retrieval, slides were pressure-cooked in 10 mM sodium citrate buffer, then cooled on ice. Endogenous peroxidase activity was quenched with 3% H_2_O_2_ for 10 min. After PBS washes, sections were blocked with PBS/0.1% Tween-20/5% normal goat serum for 1hr followed primary antibody incubations for another 1 hr at RT. Antibodies used in this study include anti-IgG (Sigma, A9044), anti-IgM (Jackson Immunoresearch, 115-035-075), anti-IgA (Bethyl Laboratories, A90-103P), anti-IgG1 (Bethyl Laboratories, A90-205P), anti-IgG2a (Bethyl Laboratories, A90-107P), anti-IgG3 (Bethyl Laboratories, A90-211P). After 3x PBS washes (10 min each), HRP activity was developed with 3,3´-diaminodbenzidine (DAB, Vector Laboratories, SK-4103), counterstained with hematoxylin, dehydrated (70%→95%→100% ethanol→xylene), and Toluene-mounted. Images were acquired using a Keyence microscope. Antibody positive area (%) was calculated as (DAB+ area within plaque intima)/(total intimal area) ×100 using Fiji (v2.1.0/1.53c).

#### **Immunofluorescence**

Following standard slide preparation (as above, omitting H₂O₂ treatment), samples were permeabilized with 0.2% Triton-X/PBS (20 min, RT) and fixed in 4% PFA (20 min). After PBS washes, sections were blocked with 5% normal goat serum in PBST (0.2% Triton-X). Primary antibodies (1:200 in blocking buffer) were incubated overnight at 4°C. IgG-AF555 (Life Technologies A31570), FcRn (abcam ab193148), CD68 (BioRad MCA1967GA), Ki67 (Abclonal A21861), NF-κB (Cell Signaling 8242), p-NF-κB (Cell Signaling 3033), F4/80 (Fisher 14-4801-81), IL-1β (Fisher P420B), ASC (Cell Signaling 67824) antibodies were used. After washing, sections were incubated with fluorescent secondary antibodies (1:400 in PBST): AF488 (Life Technologies A21206), AF568 (Life Technologies A11011), AF647 (Life Technologies A21247). Nuclei were counterstained with DAPI (1:10,000). Images were acquired using a Zeiss LSM 710 confocal microscope (20×/40×/63× objectives) or an Olympus FV3000 confocal microscope (20×/40× objectives). Positive cells were counted across multiple 63× frames (with biological replicates per condition) using Fiji (v2.1.0/1.53c).

#### **Cell culture**

RAW264.7 macrophage cells (from ATCC) were cultured in DMEM supplemented with 10% FBS and 1% Pen/Strep. Cells were passaged at approximately 70% confluency using trypsin before seeding at consistent densities across experimental plates. For induction experiments, cells underwent overnight serum starvation followed by treatment with various stimuli in media containing 0.2% fatty acid-free BSA. Treatments included: 100 μg/mL post-dialysis IgG (Cusabio CSB-NP001601m and Beyotime A7050) at time points ranging from 15 minutes to 24 hours; heat-denatured IgG (95°C for 5 minutes) at matching time points; 50 ng/mL LPS (Sigma L4524) for pro-inflammatory activation (15 minutes to 24 hours); and 1 μM TAK242 (MedChemExpress HY-11109) for 6 hours. Following treatments, cells were collected for mRNA expression analysis and WB, while centrifuged conditioned media was used for IL-1β quantification via ELISA. THP-1 cells (from ATCC) were cultured in DMEM supplemented with 10% FBS, 0.05 mM β- mercaptoethanol and 1% Pen/Strep. Cells were maintained at a cell density ranging from 5×10⁵ to 1×10⁶. Prior to experimentation, cells were induced with phorbol 12-myristate 13-acetate (PMA; 100 ng/mL) for 48 hours to differentiate monocytes into M0 macrophages. Subsequently, M0 macrophages were treated with: 100 or 200 μg/mL Human IgG (Cusabio, CSB-NP001201h), 100 μg/mL hPD1 mAb (Nivolumab, MCE HY-P9903), 1 μM TAK242, 100 ng/mL LPS (Sigma L2630), 100 nM MCC950 (NLRP3 inhibitor, MCE HY-12815), 10 μM Licochalcone D (NF-κB inhibitor, MCE HY-N4187), 10 μM Nigericin (the inflammasome signal 2 activator, MCE HY-127019) or 10 μg/mL Dil-oxLDL (Yiyuan Biotech. YB-003). Following treatments, cells and medium were collected for protein, RNA analysis, fixed cells were subjected for immunostaining analysis.

#### **Bone-marrow derived macrophages**

Femurs and tibias from 8-week-old mice were dissected, and bone marrow cells were collected in sterile PBS buffer by centrifugation. After red blood cell lysis and DMEM quenching, cells were resuspended in complete media (1 g/L glucose DMEM with 10% FBS and 1% Pen/Strep), filtered through a 100 μm strainer, and plated in 10 cm dishes. Macrophage differentiation was achieved using 50 ng/mL m-CSF (PeproTech) every 48 hours for 7 days. Differentiated BMDMs were then seeded at equal densities for experiments. For stimulation studies, serum-starved BMDMs were treated with: 100 μg/mL IgG, 20 μg/mL IgG Fab, 100 μg/mL IgG Fc, 1 μM TAK242, 50 ng/mL LPS, 100 nM MCC950, 10 μM Licochalcone D, 10 μM Nigericin, 10 μM PD98059 (ERK inhibitor; MCE HY-12028), 10 μM LY294002 (PI3K inhibitor; MCE HY-10108), 1 μM Ibrutinib (BTK inhibitor; MCE HY-10997), 50 nM Dasatinib (SRC/BCR inhibitor; MCE HY-10181), 500 nM R406 (SYK inhibitor; MCE HY-12067), 10 μg/mL Dil-oxLDL, 50 μg/mL oxLDL (Yiyuan Biotech. YB-002), 1 μM ZINC00640089 (LCN2 inhibitor, MCE HY-Q45780), or 10 μg/mL recombinant LCN2 (MCE HY-P70658). For macrophage foam cell formation, cells were incubated with 50 μg/mL oxLDL for 24 hours or with 10 μg/mL Dil-oxLDL (Yiyuan Biotech. YB-003) for 6 hours prior to harvest. For RNAi experiments, BMDMs were transfected with mouse siRNAs (RiboBio) using Lipofectamine™ RNAiMAX according to manufacturer protocols. IgG (100 μg/mL averaging 0.5 EU/mL), LPS (100 ng/mL averaging 70 EU/mL), FBS (averaging 0.01 EU/mL), and medium (averaging 0.01 EU/mL) used in this study were subjected for endotoxin tests (Beyotime, C0275S) to ensure the endotoxin-free (<0.01 EU/mL) of these reagents. Post-treatment, cells were harvested for mRNA/protein analysis, while conditioned media was centrifuged and analyzed for IL-1β secretion by ELISA.

#### **TLR4 TurboID proximity labeling**

Mouse Tlr4 cDNA sequence was synthesized by Sangon and subsequently subcloned into a pCMV-N-Flag-miniTurboID backbone (Beyotime D3034). HEK293T cells in 10 cm dishes were transfected with 10 μg of TurboID-TLR4 using PEIMAX (Polysciences 24765), together with Md2 coding plasmid. 24 hours post-transfection, the serum was replenished for 12 hours, followed by 200 μg/mL IgG treatment. Cells were incubated with 500 μM biotin for 2 hours before harvested. Biotin-labeled proteins were pulled down using Streptavidin Magnetic Beads (Invitrogen 11205D) and processed for WB analysis of TLR4-interacting proteins.

#### **Surface Plasmon Resonance**

Real-time molecular interactions were assessed using Surface Plasmon Resonance (SPR) and performed on a Biacore instrument (Cytiva Biacore 8K+) with a CM5 sensor chip^63^. Mouse recombinant TLR4 (Cusabio CSB-YP023603MO) and human recombinant TLR4-MD2 complex (R&D systems 3146-TM/CF) were diluted to 20 μg/mL with 10 M sodium acetate (pH= 4.0) and immobilized with amine coupling. Mouse and human IgG was used at the indicated concentrations and injected onto the surface of the ligand chip in 1XHBS_EP+ buffer (0.1 M HEPES; 1.5 M NaCl; 0.03 M EDTA; 0.5% P20). SPR parameters were set at 25℃, 30 μL/min flow rate, 120s contact time, and 360s disassociation time. Kinetic parameters to assess binding were analyzed using the Biacore Insight Control Software.

#### **Production of mouse Fab and Fc domains**

For preparation of IgG Fab fragments, 10-mg of mIgG (Cusabio, CSB-NP001601m) was dissolved in 100 mM phosphate buffer, pH 7.5, containing 10 mM cysteine and 2 mM EDTA. 0.25 mg of papain (Sigma-Aldrich, P4762) was subsequently added and incubated at 37°C for 3.5 hours. Proteolysis was halted by the addition of 1 mL of 20 mM iodoacetamide in the same buffer to inhibit an enzymatic reaction. The resulting product was analyzed by SDS-PAGE to verify the presence of both Fc and Fab fragments. A column packed with Recombinant Protein A+G (Beyotime, P2019) was first equilibrated with PBS. Papain-treated mIgG was dialyzed in the PBS. This solution was then loaded onto the pre-equilibrated sorbent bed, and PBS was used to wash the column. Flow-through was collected, containing IgG Fab fragments. The coding DNA sequence for mouse IgG2a-Fc (E215–K447) was cloned into a pcDNA3.4 vector. Vectors were then transfected in 293-F cells. 7 days later, the supernatant was collected and purified using a Recombinant Protein A+G column according to manufacturer instructions. Protein solution was concentrated using a displaced buffer with 3-kD cut-off Amicon (Millipore).

#### **Quantitative real-time PCR**

Total RNA was extracted from tissues or cells using the Tri-Isolate RNA Pure Kit (IBI Scientific IB47632). cDNA synthesis was performed with the High-Capacity cDNA Reverse Transcription Kit (Thermo Fisher Scientific). Quantitative PCR was conducted using AzuraView GreenFast qPCR Blue Mix on a Bio-Rad CFX96 system, with gene expression normalized to *Hprt* (BMDMs/RAW264.7 cells) using the ΔΔCt method. QPCR primer sequences are available upon request.

#### **Western blots**

Protein lysates from aortic arches or cells were prepared using the IntactProtein™ Cell-Tissue Lysis Kit (GenuIN Biotech #415) supplemented with protease inhibitors. Tissues were homogenized immediately after dissection using a Polytron homogenizer, with all procedures performed on ice. Protein concentrations were determined by BCA assay (ThermoFisher 23225). For plasma samples, sample was diluted 1:100 in ddH₂O prior to SDS-PAGE. WB was performed using the following primary antibodies: Anti-IgG (Sigma A9044), Anti-IgM (Jackson ImmunoResearch 115-035-075), Anti-IgA (Bethyl A90-103P), Anti-HSP90 (Proteintech 13171-1-AP), Anti-phospho-NF-κB (Cell Signaling 3033T), Anti-NF-κB (Cell Signaling 8242T), Anti-GAPDH (Cell Signaling 14C10), Anti-NLRP3 (Proteintech 30109-1-AP), Anti-pro-IL-1β (Proteintech 26048-1-AP), Anti-pro-IL-18 (Cell Signaling 57058), Anti-Caspase 1/p20 (Proteintech 22915-1-AP), Anti-LCN2 (Proteintech 26991-1-AP). Proteins were detected by enhanced chemiluminescence (ECL) (ThermoFisher 32106). Coomassie Brilliant Blue gel staining provided loading control bands for plasma samples.

#### **RNA sequencing and analysis**

RNA was extracted from IgG-treated BMDMs and subjected to sequencing. Libraries were prepared and sequenced on an Illumina NovaSeq 6000 platform (Novogene). Quality control was performed using FastQC (v0.11.9), followed by alignment to the GRCm39 mouse genome with STAR (v2.7.7a). Gene counts were generated using featureCounts (v2.0.1) and normalized via the TMM method. Differential expression analysis compared IgG-treated versus control cells using log2(FPKM+0.5) values, with DEGs defined as |log2FC| > 0.75 and adjusted p-value < 0.1. Results were visualized as Z-score heatmaps. The dataset is publicly available in the Genome Sequence Archive (PRJCA020637).

**SUPPLEMENTAL FIGURES**

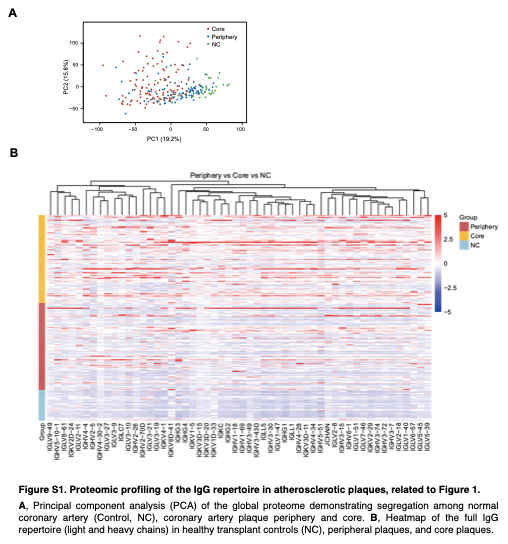

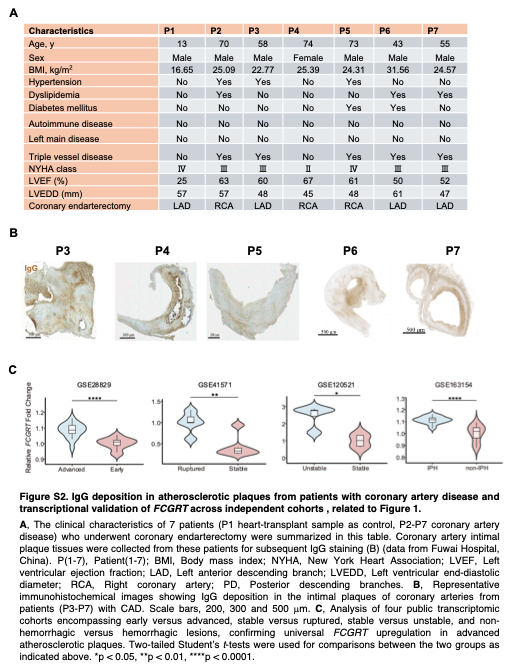

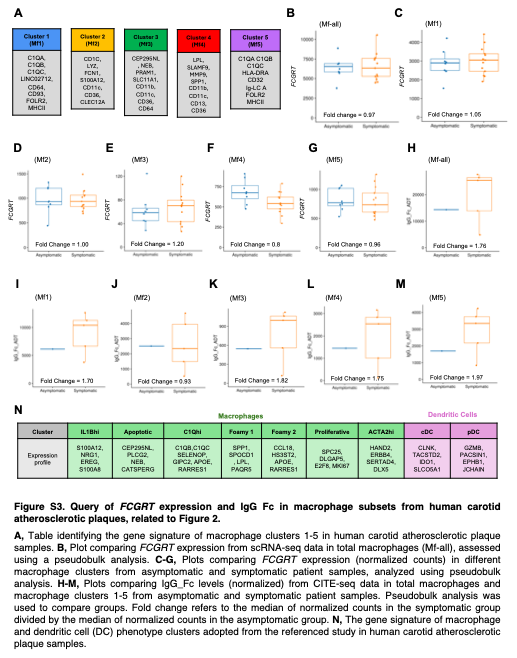

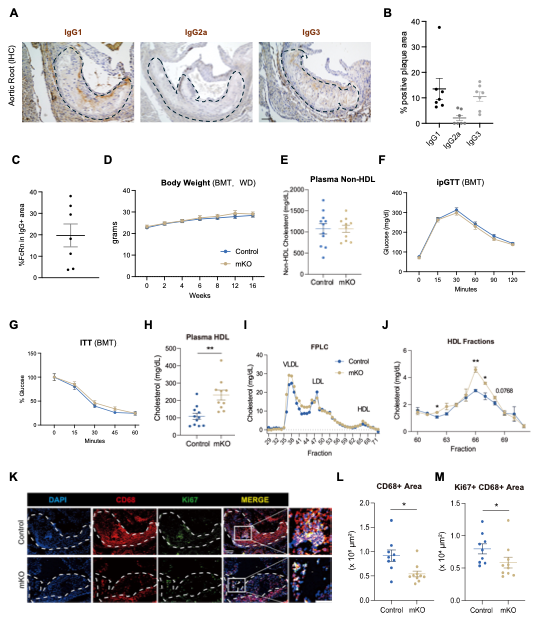

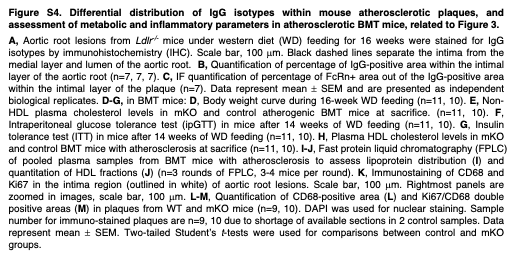

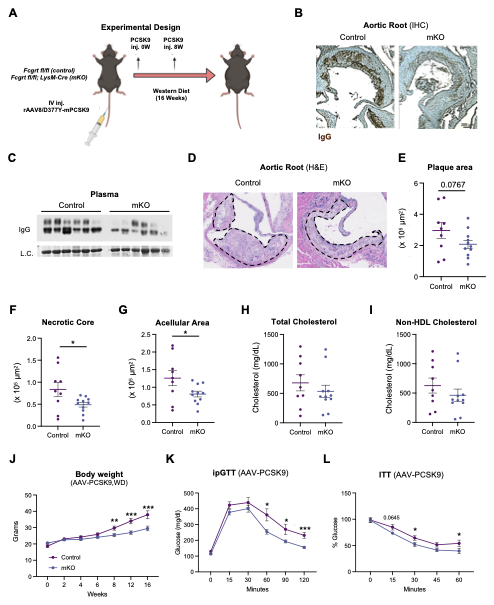

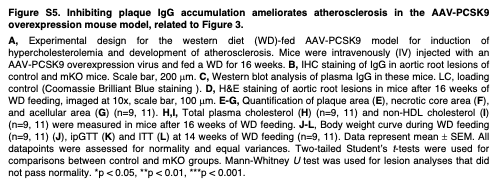

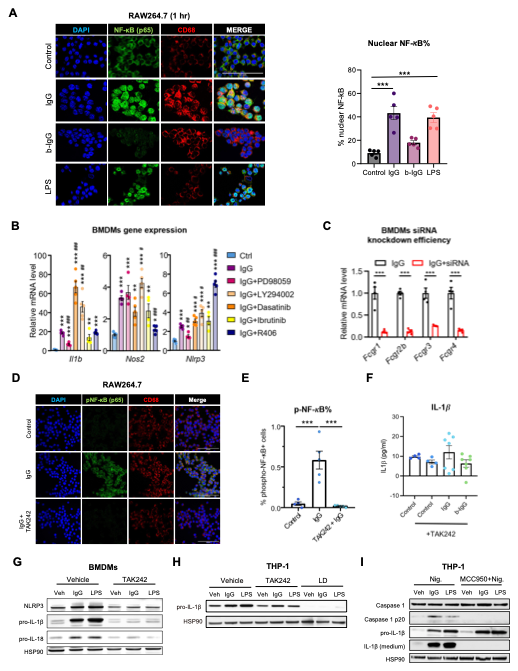

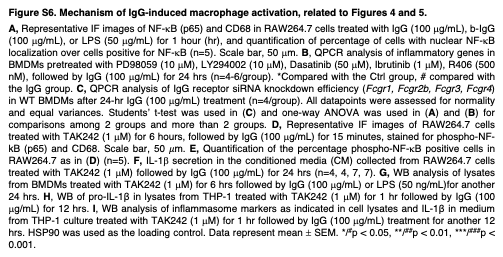

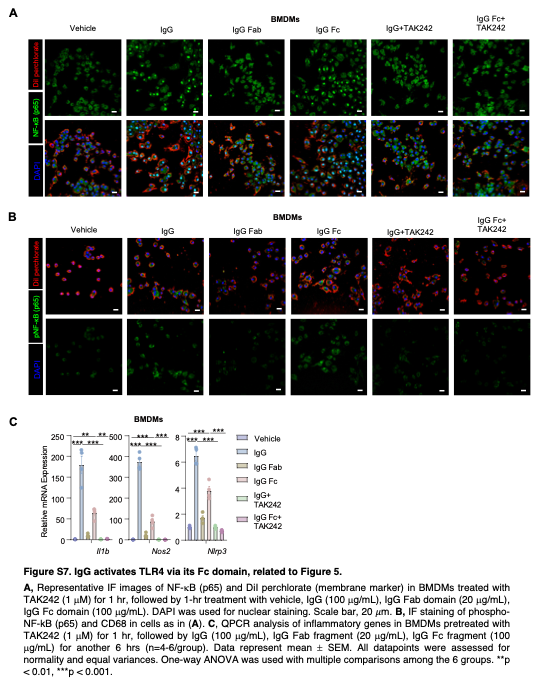

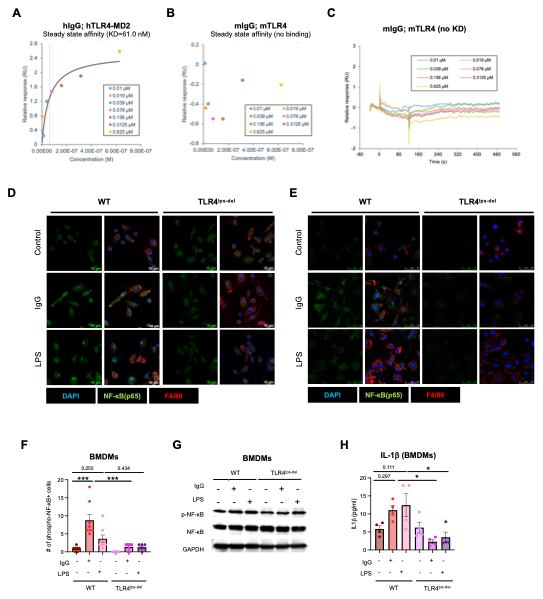

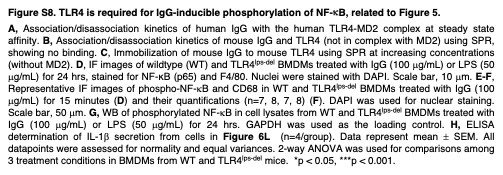
